# A therapeutic hepatitis B mRNA vaccine with strong immunogenicity and persistent virological suppression

**DOI:** 10.1101/2022.11.18.517095

**Authors:** Huajun Zhao, Xianyu Shao, Yating Yu, Lulu Huang, Narh Philip Amor, Kun Guo, Changzhen Weng, Weijun Zhao, Ailu Yang, Jiesen Hu, Hongbao Yang, Zhenguang Liu, Qiuju Han, Leilei Shi, Shiyu Sun, Jian Zhang, Yong Yang, Ang Lin

## Abstract

Here we report on the development and comprehensive evaluations of an mRNA vaccine for chronic hepatitis B (CHB) treatment. In two different HBV carrier mouse models generated by viral vector-mediated HBV transfection (pAAV-HBV1.2 and rAAV8-HBV1.3), this vaccine demonstrates sufficient and persistent virological suppression, and robust immunogenicity in terms of induction of strong innate immune activation, high-level virus-specific antibodies, memory B cells and T cells. mRNA platform therefore holds prospects for therapeutic vaccine development to combat CHB.

## INTRODUCTION

CHB is one major cause of liver fibrosis, cirrhosis, and hepatocellular carcinoma and poses a major public health threat^1^. Current CHB treatments such as nucleoside/nucleotide- and interferon-alpha (IFN-α)-based therapy, were unable to achieve sufficient viral clearance^2,3^. Multiple approaches to CHB therapeutic vaccines have been practiced intensively, including recombinant protein-based subunit, adenoviral vectored, and DNA-based vaccines, but yet achieved efficient seroclearance of HBsAg and seroconversion of anti-HBs antibody (Ab)^4,5,6^. mRNA vaccines that contain antigen-encoding mRNAs encapsulated into for example lipid nanoparticle (LNP) have shown superior immunogenicity in eliciting both Ab and cellular immune responses over other types of vaccines, and are also endowed with strong intrinsic adjuvant property to activate innate immune compartment^7^. Several mRNA-based prophylactic or therapeutic vaccines for infections, malignancies or other diseases are currently being evaluated in clinical trials ^8, 9, 10, 11^. This platform also holds prospects for the development of therapeutic CHB vaccine that is expected to elicit potent anti-viral immunity mediating efficient virological suppression.

## RESUTLS AND DISCUSSION

Recently, we reported a proprietary artificial intelligence-based algorithm that designs mRNA with optimal folding stability and codon usage that together contribute to a high translation efficiency ^12^. Using this algorithm, a novel mRNA vaccine for CHB treatment was developed, which is composed of Hepatitis B surface antigen (HBsAg)-encoding mRNAs encapsulated into an ionizable lipid-based LNP through a well-established microfluidic system^13^. The mRNAs were modified with N1-Methyl-pseudouridine and showed efficient protein expression reaching a mean level of 980.6 mIU/ml upon transfection into HEK-293T cells (Supplemental Figure 1A). Vaccine formulations were well characterized which demonstrated particle diameters of 96.3 ± 2.16 nm with average zeta potential of −1.92 mV and polymer dispersity index (PDI) below 0.2. In addition, cells incubated with escalating concentrations of mRNA vaccines for 24 hours maintained high viability, which suggested a limited cytotoxicity of vaccine (Supplemental Figure 1B).

Therapeutic efficacy of the mRNA vaccine was first evaluated in pAAV-HBV1.2-transduced HBV-carrier mice. This model shows systemic immune tolerance and long-lasting HBV viremia that largely resemble asymptomatic chronic HBV-infected individuals and has therefore been widely used in the study of CHB immunotherapy^14,15^. HBV-carrier mice were administered intramuscularly (i.m.) with three doses of 5μg or 10μg mRNA vaccines at a 1-week interval (Figure 1A). Compared to PBS-treated mice showing a high serum HBsAg level, mRNA vaccine-treated mice demonstrated a rapid decline of serum HBsAg, which was even undetectable 7 days after the 3^rd^ dose (Figure 1B). Clinical management of CHB remains challenging largely due to HBV recurrence and the failure to achieve seroconversion of anti-HBs Abs^2, 3^. Magnitude of anti-HBs Ab response was therefore evaluated longitudinally. Three doses of HBV mRNA vaccines elicited robust levels of anti-HBs Abs reaching a mean titer of 3624.0 mIU/ml and 4804.6 mIU/ml in the 5μg and 10μg vaccine groups at day 59, respectively (Figure 1C).

**Figure 1.**
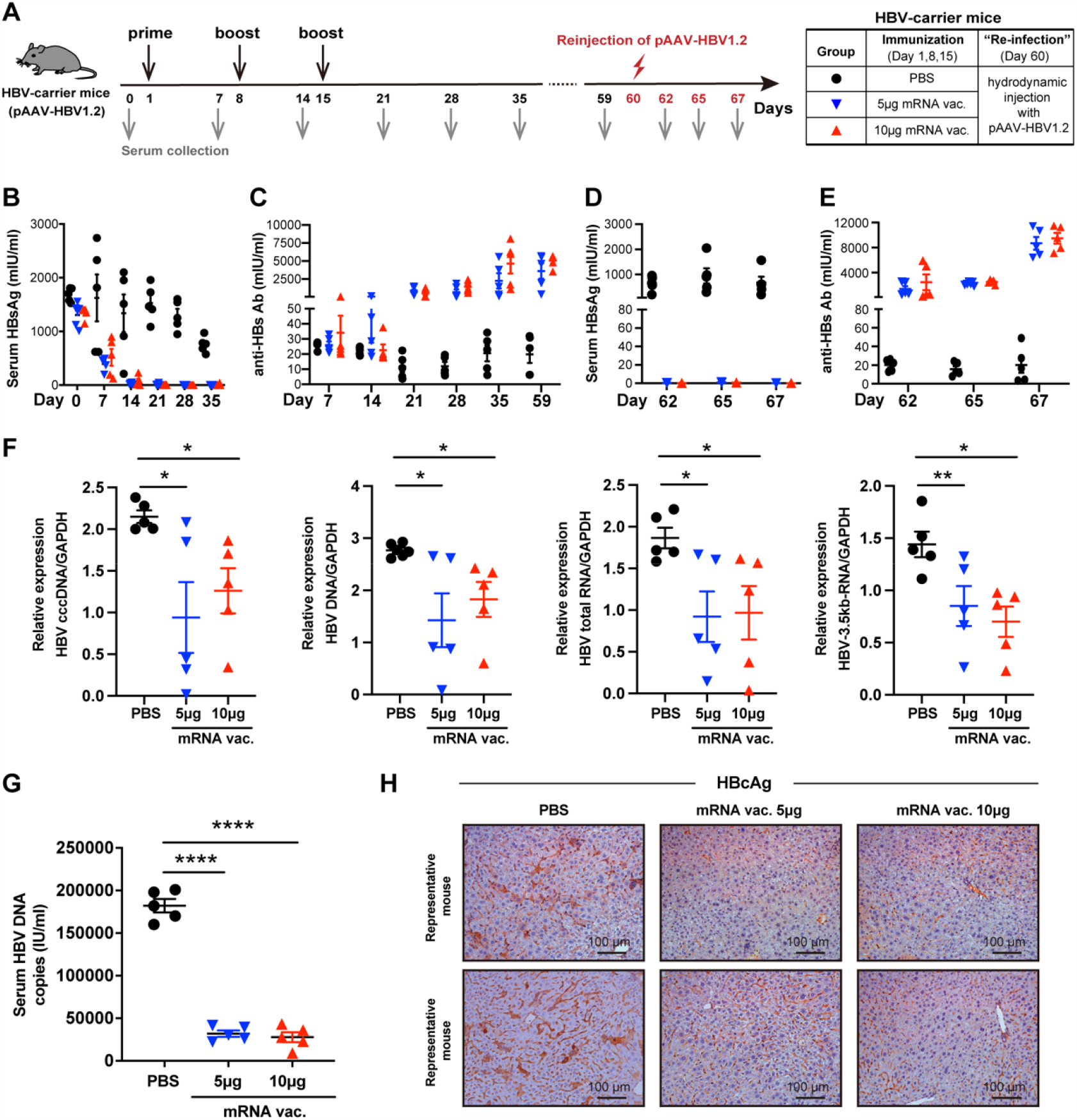
HBV mRNA vaccine induced efficient viral suppression and achieved robust seroconversion in pAAV-HBV1.2 mice. (A) Study design. pAAV-HBV1.2-transduced HBV-carrier mice (n=5/group) were immunized i.m. with 5μg or 10μg HBV mRNA vaccines three times at a 1-week interval. HBV-carrier mice administered with PBS were used as control. At day 60, mice were hydrodynamically re-injected with 8μg pAAV-HBV1.2 plasmids. Sera samples were collected at the indicated time points. (B, D) Levels of serum HBsAg and (C, E) anti-HBs Abs were measured at the indicated time points. (F) Seven days after viral re-exposure, intrahepatic HBV cccDNA, total DNA, total RNA and 3.5-kb RNA were analyzed. (G) Seven days after viral re-exposure, serum HBV DNA was quantified. (H) Seven days after viral re-exposure, HBcAg expression in liver tissues was determined by IHC staining (magnification: ×200; scale bar: 100 μm). GAPDH, glyceraldehyde 3-phosphate dehydrogenase; SEM, standard error of the mean. Data are shown as Mean ± SEM. ^*^*p* ≤ 0.05, ^**^*p* ≤ 0.01, ^****^*p* ≤ 0.0001.

To evaluate whether the mRNA vaccine could induce sustained protective responses against viral re-exposure, the mice were re-injected with pAAV-HBV1.2 plasmids on day 60 mimicking viral challenge. As expected, PBS-treated mice still demonstrated a high level of serum HBsAg. While all vaccinated mice were fully protected against viral re-exposure showing no detectable level of serum HBsAg (Figure 1D), which was accompanied by further elevated anti-HBs Ab titers (Figure 1E). Moreover, copies of intrahepatic HBV cccDNA, total DNA, total RNA, 3.5kb RNA (Figure 1F) and serum HBV DNA (Figure 1G) were clearly reduced in vaccinated mice and the expression of HBV core antigen (HBcAg) in liver tissues was largely decreased (Figure 1H). Since AAV-transduced-HBV-carrier mice cannot produce significant amounts of cccDNA, the anti-HBV effect on eliminating HBV cccDNA should be further evaluated in specific animal models, such as rc-cccDNA mouse model^16^ or human liver chimeric mouse model^17^.

Efficacy of the mRNA vaccine was next compared side-by-side with conventional therapeutic Entecavir. Compared to the rapid anti-HBs Ab production and serum HBsAg clearance induced by mRNA vaccine, Entecavir administered orally for consecutive 15 days showed no ability in inducing anti-HBs Abs or eliminating serum HBsAg (Supplemental Figure 2), which was in line with previous studies^18^. Considering that viral clearance can be mediated through vaccine-induced cytotoxic immune responses potentially causing liver damage, serum alanine transaminase (ALT) and aspartate aminotransferase (AST) levels were monitored longitudinally (Supplemental Figure 3A-B). A transient elevation of serum ALT was observed in a few animals 7 days post the 1^st^ and 2^nd^ vaccination. While serum AST remained at normal levels in all treated mice during the period of treatment. In addition, histopathological analysis revealed that there was no hepatotoxicity or liver injury induced upon the 3-dose vaccine treatment (Supplemental Figure 3C). These suggested that the mRNA vaccine may potentially function through a non-cytotoxic mechanism to eliminate virus. Previously, we and others showed that IFN-γ-producing HBV-specific CD8+ T cells could mediate HBV clearance via a non-cytotoxic mechanism without causing liver damage^19,20^. Since both cytotoxic and non-cytotoxic immune responses to HBV are important to eliminate virus, the underlying mechanisms of action of the mRNA vaccine would merit further investigation.

Therapeutic efficacy of the mRNA vaccine was further evaluated in rAAV8-HBV1.3-transduced HBV-carrier mouse model (Figure 2A), which shows more efficient and homogeneous HBV transduction than the aforementioned pAAV-HBV1.2 mice model^21^. As validated in our study, rAAV8-HBV1.3-transduced mice showed persistent HBV surface antigenemia lasting for more than 200 days (Supplemental Figure 4) and a typical immunotolerant status represented by high frequencies of hepatic CD4^+^CD25^+^ Foxp3^+^ T cells (Supplemental Figure 4B), elevated expression of inhibitory immune-checkpoint molecules (PD-1, LAG-3, and TIM-3), and impaired cytokine production by CD4^+^ T and CD8^+^ T cells upon stimulation (Supplemental Figure 4C and D). All these features well mimic the immunotolerant state of human chronic HBV carriers^21^. Upon three doses of vaccination, HBV1.3-carrier mice demonstrated a rapid and sufficient serum HBsAg clearance, and the virological suppression was maintained for at least 208 days within the period of observation (Figure 2B). Furthermore, serum HBeAg level was remarkably reduced in both two groups of vaccinated mice (Figure 2C). Serum HBV DNA copies were significantly reduced in mice receiving 10μg mRNA vaccine and showed a trend of decrease in the 5μg dosing group (Figure 2D). Notably, majority of the 10μg mRNA vaccine-treated mice still presented high levels of anti-HBs Abs when detected 208 days after treatment initiation (Figure 2E). Since serum HBsAg was sufficiently cleared, but serum HBV DNA was still present which implied that viral replication may still remain active at a very low level although was largely restricted. The mRNA vaccine-elicited anti-viral immunity remains to be further assessed for the efficacy to control viral genome transcription using more suitable animal models.

**Figure 2.**
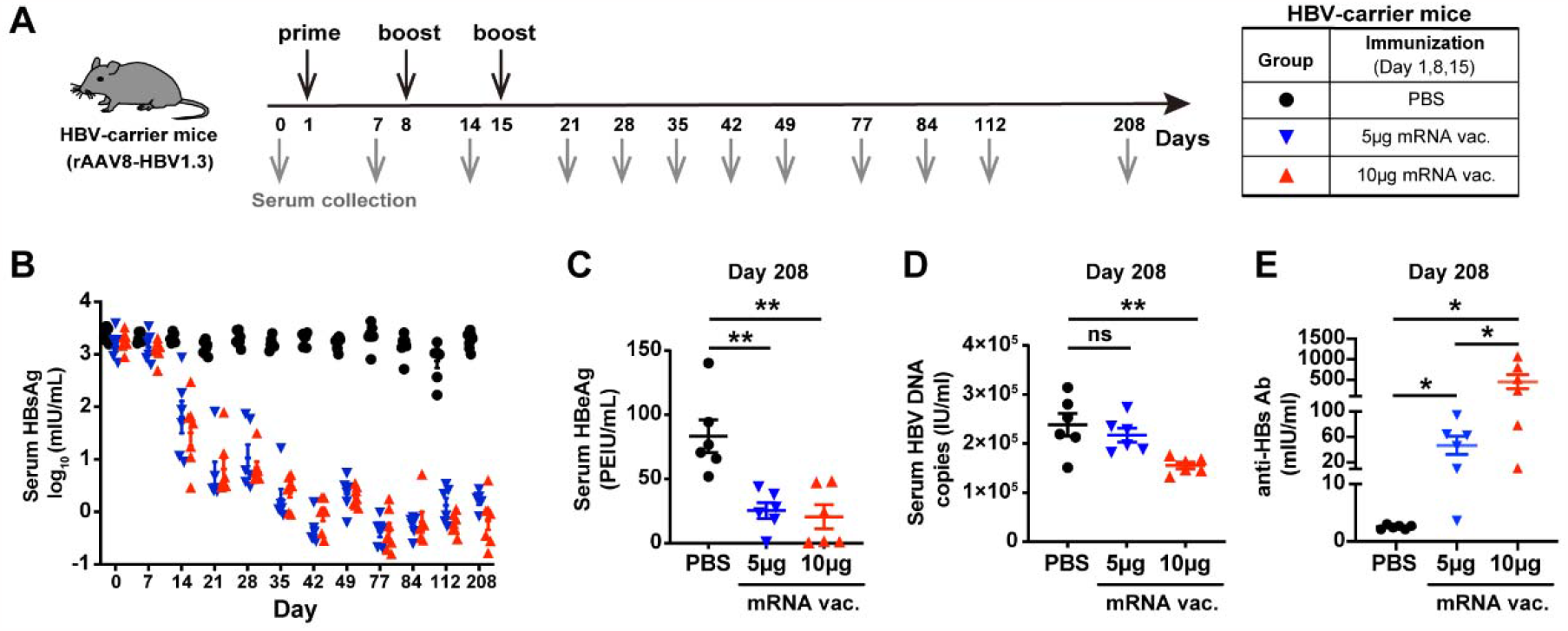
HBV mRNA vaccine induced efficient and sustained viral suppression and achieved robust seroconversion in rAAV8-HBV1.3 mice. (A) Study design. rAAV-HBV1.3-transduced HBV-carrier mice (n=6/group) were immunized i.m. with 5μg or 10μg HBV mRNA vaccines three times at a 1-week interval. HBV-carrier mice administered with PBS were used as control. Sera samples were collected longitudinally. (B) Levels of serum HBsAg were measured at the indicated time points. (C-E) 208 days after treatment start, levels of serum HBeAg (C), serum HBV DNA copies (D), and anti-HBs Abs (E) were measured. Data are shown as Mean ± SEM. ^*^*p* ≤ 0.05, ^**^*p* ≤ 0.01. PEIU represents Paul-Ehrlich-Institute Unit.

To further validate the potency and superiority of the mRNA vaccine, we next performed a side-by-side comparison with our recently reported recombinant CHB therapeutic vaccines (Sim+rHBV)^22^ in rAAV8-HBV1.3 mice. Three doses of Sim+rHBV vaccine showed a moderate ability in viral clearance but much less efficient than the mRNA vaccine (Supplemental Figure 5). Seven days post the 3^rd^ dose, serum HBsAg levels in mice receiving 5μg mRNA vaccine, 10μg mRNA vaccine and Sim+rHBV vaccine were present at a mean value of 15.8 mIU/ml, 16.0 mIU/ml and 166.0 mIU/ml, respectively. In addition, evidence from other groups testing their in-house developed therapeutic CHB vaccines using same animal models indirectly supported the superior therapeutic effect of mRNA vaccine^23, 24^.

An optimal therapeutic vaccine for CHB treatment should have the potentials to trigger efficient innate immune activation leading to robust antigen presentation and the resulting generation of HBV-specific cellular immunity^12^. To this end, we studied the innate immune responses induced at early time point (12 hours) after prime immunization in rAAV8-HBV1.3 mice and observed an increased infiltration of dendritic cell (DC) subsets (CD8α^+^ cDC1s, CD103^+^ cDC1s and CD11b^+^ cDC2s) and macrophages into spleen, accompanied by potent cell maturation (Supplemental Figure 6A-D). Other innate immune cell subsets including monocytes and neutrophils (Supplemental Figure 6E-G) also showed a phenotypic maturation, indicated by elevated expression of CD80 and CD86. T cell exhaustion is a hallmark of CHB infection and the restored HBV-specific CD8^+^ T cell function has been widely-accepted to be predictive for the efficacy of therapeutic vaccines for CHB^25^. We therefore assessed whether the therapeutic mRNA vaccination could promote HBV-specific T cell responses and break the exhaustion. In rAAV8-HBV1.3 mice, three doses of mRNA vaccines induced robust levels of Th1-biased CD4^+^ and CD8^+^ T cells producing IFN-γ or IL-2 favoring viral elimination (Supplemental Figure 7A-C). In addition, frequencies of HBsAg-specific memory B cells (MBCs) in spleens were obviously elevated upon vaccination (Supplemental Figure 7D) which was associated with the strong seroconversion and long-term protection as observed earlier in this study (Supplemental Figure 2).

Collectively, we reported an HBV mRNA vaccine candidate with potent therapeutic efficacy and strong immunogenicity. To our knowledge, this is the first reported therapeutic mRNA vaccine candidate for CHB treatment. Three doses of HBV mRNA vaccines could efficiently and persistently eliminate HBV and achieve a long-term seroconversion of anti-HBs Ab, and most importantly showed full protection against subsequent viral re-exposure. The strong innate immune activation and generation of robust functional HBV-specific T cells and MBCs by the mRNA vaccine may hold prospects for functional cure of CHB and prevention of HBV recurrence. However, further in-depth assessment of the mRNA vaccine would be needed to evaluate the effects on restricting viral replication at genomic levels especially the potentiality to eliminate HBV cccDNA pool. In addition, synergistic efficacy of the mRNA vaccine in combinatorial use with other types of CHB therapeutics merits further investigation.

## METHODS

A detailed description and additional materials and methods are available in supplemental materials.

### Ethics, animals, treatments

C57BL/6J mice (5-6 weeks old, male) were purchased from Beijing HFK Bioscience Co. Ltd. (Beijing, China). HBV-carrier mouse models were generated either through hydrodynamic injection of a volume of saline (equivalent to 10% body weight) containing 8μg pAAV-HBV 1.2 plasmid (kindly provided by Pei-Jer Chen; National Taiwan University College of Medicine, Taipei, Taiwan) or intravenous injection of 1 × 10^10^ vector genome equivalent of rAAV8-HBV1.3, as previously described^21,23^. Serum HBsAg levels were measured 6 weeks after the hydrodynamic or intravenous injection, and mice with serum HBsAg levels > 500 mIU/mL were defined as HBV-carrier mice and used for subsequent in-vivo experiments. HBV-carrier mice were randomly allocated to different groups and were i.m. injected with three doses of 5μg or 10μg mRNA vaccines at a one-week interval. HBV-carrier mice administered with PBS were used as control. In some experiments, HBV-carrier mice were orally treated with Entecavir (50μg/kg, Selleck Chem) for 15 days consecutively or were immunized with a recombinant therapeutic vaccine (Sim+rHBV)^21^ containing 2 μg rHBVvac adjuvanted with 100 μg simvastatin. Sera samples were collected at different time points and were stored at −80 ºC for further use. For the evaluation of long-term protective response, the immunized HBV-carrier mice were hydrodynamically re-injected with 8μg pAAV-HBV1.2 plasmids on day 60 after treatment initiation. All animal experiments were performed in accordance with the Guidelines for the Care and Use of Laboratory Animals and the Ethical Committee of Shandong University and using protocols approved by the Institutional Animal Care and Use Committee of Shandong University (approval number: 20023).

### mRNA vaccine preparation

mRNAs encoding for HBsAg (Genbank accession number: MK213856.1) were synthesized by T7 polymerase-mediated in vitro transcription (IVT) based on a linearized DNA template containing codon-optimized HBsAg gene flanked with 5’ and 3’ untranslated regions (UTRs) and a 100 nt poly-A tail. During IVT procedure, mRNAs were modified with N1-Methyl-pseudouridine (Synthgene) and capped using CleanCap Reagent (TriLink). After this, IVT products were purified with Monarch RNA purification columns (NEW ENGLAND BioLabs Inc. MA, USA) and resuspended in a TE buffer at a desired concentration. For mRNA encapsulation into LNP, lipid components were dissolved in ethanol at molar ratios of 50:10:38.5:1.5 (ionizable lipid: DSPC: cholesterol: DMG-PEG2000). The novel ionizable lipid (YX-02) was designed and has been patented by Firestone Biotechnologies. The lipid cocktail was mixed with mRNAs dissolved in 10mM citrate buffer (pH4.0) at an N/P ratio of 5.3 :1 and a volume ratio of 3: 1 using a microfluidic-based equipment (INano™L from Micro&Nano Biologics) at a total flow rate of 12 mL/min. Formulations were diluted with PBS and ultrafiltrated using 50-kDa Amicon ultracentrifugal filters. Vaccine formulation was characterized for particle diameter, polymer dispersity index (PDI) and zeta potentials using NanoBrook Omni ZetaPlus (Brookhaven Instruments).

## STATISTICAL ANALYSIS

Statistical analyses were performed using GraphPad Prism software (v6.0; GraphPad Software, La Jolla, CA, USA). An unpaired Student’s *t* test was applied for comparisons between two groups, and differences among multiple groups were analyzed by one-way analysis of variance. A *p* value less than 0.05 was considered statistically significant (^*^p ≤0.05, ^**^p ≤ 0.01, ^***^p ≤ 0.001).

## DATA AVAILABILITY

All data are available upon reasonable request to the corresponding authors.

### ACKNOWLEDGEMENTS

This work was supported by the Natural Science Foundation of Jiangsu Province (BK20221031 to A.L.), the National Natural Science Foundation of China (32200764, 82061138008), the National Postdoctoral Program for Innovative Talents (BX20190192, to J.H.Z.), the Fundamental Research Funds for the Central Universities (2632022YC01, to A.L.), the National Key Research and Development Program (2021YFC2300603, to J.Z.), the National Science Foundation for Young Scientists of China (82001687, to J.H.Z.), the Shandong Provincial Natural Science Foundation for The Excellent Youth Scholars (ZR2022YQ75, to J.Q.H), and the National Natural Science Foundation of China (81972686, to J.Q.H). This study was also supported by the Projects of State Key Laboratory of Natural Medicines (SKLNMZZ2023XX) and State Key Laboratory of Microbial Technology (M2023-13). We thank the Pharmaceutical Biology Sharing Platform and the Translational Medicine Core Facility of Shandong University and China Pharmaceutical University for instrumental support. We also thank Firestone Biotechnology Co., Ltd for technical assistance with the development of novel ionizable lipids and thank Dr. Liang Zhang for the assistance in codon optimization of mRNA using LinearDesign Algorithm.

## AUTHOR CONTRIBUTIONS

A.L., H.Z., Y.Y. and J.Z. designed the project; H.Z., X.S., Y.Y., L.H., N.A., C.W., W.Z., A.Y., J.H., Q.H., L.S. and S.S. performed experiments; H.Z., A.L., X.S., T.Y., B.Y. and N.A. performed data analysis; A.L., Y.Y., and H.Z. discussed the data; A.L. H.Z., Y.S. and L.H. wrote the manuscript.

