## Supplemental Material for "A therapeutic hepatitis B mRNA vaccine with strong immunogenicity and persistent virological suppression"

**Supplemental Materials and Methods**

**Evaluation of mRNA translation in vitro**

Human embryonic kidney (HEK) 293T cells were cultured in high-glucose Dulbecco's Modified Eagle Medium (DMEM, BIOIND) supplemented with 10% fetal bovine serum (FBS, BIOIND) and 1% penicillin-streptomycin (NCM Biotech). 5 × 10^5^ HEK293T cells were seeded into 6-well plates and were transfected with 2μg HBsAg-encoding mRNAs using jetMessegner® transfection reagent (Polyplus-transfection^®^) according to the instructions. Cells were harvested 48 hours later and were lysed using RIPA lysis buffer (Beyotime). Levels of HBsAg in cell lysates were quantified using HBsAg Quantification Kit (Autobio).

**Cytotoxicity of HBV mRNA vaccine**

HEK-293T or AML12 cells were seeded into 96-well plates at a density of 8×10^3^ cells per well suspended in 100 μL DMEM medium. Cells were incubated with or without escalating concentrations of HBV mRNA vaccines (50μg/ml, 100μg/ml, 200μg/ml, 400μg/ml, 800μg/ml, 1600μg/ml, 3200μg/ml) for 24 hours. Cell Counting Kit-8 (CCK-8) was used to determine cytotoxicity according to the manuals. Briefly, CCK-8 reagent was added into cell culture prior to incubation at 37 ℃ for 1 hour. After this, absorbance of mixture was read at 450 nm wavelength using CMax Plus microplate reader (Molecular Device). Cell viability was calculated using formula: Cell viability (%) = (OD1-OD3) / (OD2-OD3) × 100%. OD1: cells treated with mRNA vaccines; OD2: cells cultured with medium alone; OD3: DMEM medium only.

**Quantification of serum HBsAg and HBeAg**

Levels of serum HBsAg and HBeAg were quantified using Chemiluminescence Immunoassay (CLIA) Commercialized Kits according to the manuals (Autobio). Briefly, undiluted sera (50μL) were added into the wells followed by incubation with detection reagent (50μL) at 37 ℃ for 60 minutes. After washing, chemiluminescent substrates (50μL) were added into the mixture and incubated at room temperature (RT) for 10 minutes in the dark. Plates were read using Synergy 2 Multi-Mode Microplate Reader (BioTek, Vermont).

**Determination of anti-HBs IgG titer**

Titers of serum anti-HBs Abs were determined using commercialized ELISA kit (Wantai Bio-pharm) according to instruction. Briefly, undiluted sera (50μL) were added into wells pre-coated with antigens followed by incubation with horse radish peroxidase (HRP)-conjugated anti-mouse IgG at 37 ℃ for 60 min. After washing, TMB substrate was used for development and the absorbance was read at 450 nm (minus 630 nm for wavelength correction) using the Synergy 2 Multi-Mode Microplate Reader (BioTek, Vermont).

**Measurement of serum ALT and AST**

Levels of serum ALT and AST were quantified using commercialized ELISA kits according to the manuals (Nanjing Jiancheng Bioenginering Institute). Plates were read using Synergy 2 Multi-Mode Microplate Reader (BioTek, Vermont).

**Histopathological and immunohistochemical analysis**

Liver tissues were collected and fixed in 4% paraformaldehyde (Sinopharm Chemical Reagent) and embedded in paraffin wax. 5-μm tissue sections were cut, dewaxed and rehydrated through xylene and alcohols and were subsequently subjected to hematoxylin and eosin (H&E) staining for histopathological assessment as previously described^1^. For Immunohistochemical (IHC) staining, tissue sections were dewaxed, rehydrated and antigen retrieved using proteinase K antigen retrieval solution (Abcam). Following this, tissue sections were washed three times with Tris-buffered saline (TBS) buffer and incubated with goat anti-Rat IgG (OriGene) as blocking reagent for 15 min. Intrahepatic expression of core antigen of HBV (HBc) was detected by incubating with anti-HBcAg monoclonal Ab (Gene Tech; GB058629) overnight at 4 ℃. After washing with TBS buffer, tissue sections were incubated with biotinylated anti-rabbit IgG and streptavidin/horseradish peroxidase conjugates (ZSGB-Bio) for 20 minutes at 37 ℃. DAB substrate was then added followed by counterstaining with hematoxylin. Finally, sections were dehydrated, cleared and mounted for analysis. Images were captured using Olympus BX46 microscope (Olympus, Tokyo, Japan).

**Quantification of HBV DNA and HBV RNA**

Serum HBV DNA was measured by qPCR using an HBV DNA kit (Sansure Biotech). Intrahepatic HBV genomic DNA was extracted using a genomic DNA kit (Tiangen Biotech). Intrahepatic total RNA was extracted using TRIzol reagent (Invitrogen), and the RNA was reverse-transcribed into cDNA using a commercially available cDNA synthesis kit (CW Biotech). To distinguish the HBV cccDNA from the pAAV-HBV episome (plasmid DNA), the genomic DNA were treated by multi-enzyme digestion based the unique restriction site (SwaI) of pAAV-HBV1.2 plasmid as previously described^2^. Briefly, 1μg of extracted DNA was digested with 10 U of SwaI restriction enzyme (New England Biolabs) for 15 min at 25°C and then 1 U of ATP Dependent DNase (Takara Biomedical Technology) was added followed by incubation at 37°C for 16 hours. Real-time PCR for intrahepatic HBV DNA and RNA was performed by a Lightcycler^®^ 96 system (Roche) using the UltraSYBR mixture (CW Biotech). Sequences of the primers used are listed in Supplemental Table 1.

**Isolation of splenic mononuclear cells**

Splenic mononuclear cells (MNCs) were isolated as previously described^2,3^. Briefly, spleen tissues were grinded gently and washed through a 200-μm sterile cell strainer. Cells were re-suspended in PBS and centrifuged at 400 *g* for 10 minutes. Cell pellets were next re-suspended with a suitable volume of Red Blood Cells (RBC) lysis buffer (Solarbio) for 5 minutes at 4 ℃. Following this, 1× PBS was added to stop the RBC lysis and centrifuged at 400 g for 10 minutes to obtain splenic MNCs. After centrifugation, cells were re-suspended in RPMI-1640 medium containing 10% fetal bovine serum (FBS, BIOIND) and 1% penicillin-streptomycin (NCM Biotech) for subsequent in-vitro experiments.

**Isolation of hepatic mononuclear cells**

Hepatic mononuclear cells (MNCs) were isolated as previously described^2,3^. Liver tissues were grinded gently and washed through a 200-μm sterile cell strainer. Cell suspensions were first centrifuged at 100g for 1 minute to precipitate and remove hepatocytes. Subsequently, cell supernatants were centrifuged at 400 g for 10 minutes and cell pellets were resuspended in 40% Percoll (GE Healthcare) followed by centrifugation at 800g for 25 minutes. Cell pellets were next re-suspended with a suitable volume of Red Blood Cells (RBC) lysis buffer (Solarbio) for 5 minutes at 4 ℃. Following this, 1× PBS was added to stop the RBC lysis and centrifuged at 400 g for 10 minutes to obtain hepatic MNCs. After centrifugation, cells were re-suspended in RPMI-1640 medium containing 10% fetal bovine serum (FBS, BIOIND) and 1% penicillin-streptomycin (NCM Biotech) for subsequent in-vitro experiments.

**Characterization of rAAV8-HBV1.3-transduced HBV-carrier mice**

rAAV8-HBV1.3-transduced HBV-carrier mice were characterized by evaluating several aspects of T cell function as demonstrated in Supplemental Figure 4. For the evaluation of cytokine-producing ability of T cells, 2 million hepatic MNCs were seeded per well into 96-well plates and incubated with 30ng/mL PMA and 1μg/mL ionomycin (both ordered from Beyotime) for 4 hours in the presence of 5μg/mL brefeldin A (Biolegend). Cytokine production of T cells was evaluated by surface and intracellular staining using Fixation/Permeabilization Solution Kit (BD Biosciences) according to the manuals. For the analysis of PD-1, LAG-3, and TIM-3 expression on T cells, hepatic and splenic MNCs were used and first incubated with Live/Dead Fixable Blue Dead Cells Stain Kit (Thermofisher) for 5 minutes. After washing, cells were incubated with antibody cocktails for 20 minutes at 4 ℃ in dark. Flow cytometry analysis was carried out on BD FACSymphony A3 (BD Biosciences). Data was analyzed using FlowJo V10.8 software (Tree Star). Fluorochrome-conjugated antibodies used in this study are listed in Supplemental Table 2.

**Analysis of antigen-specific memory B cell (MBC) response**

Frequencies of HBsAg-specific MBCs were assessed by flow cytometry. For the preparation of HBsAg probes, HBsAg was first biotinylated using a Biotin Quick Labeling Kit (Friendbio Science) according to the manuals. Biotinylated HBsAg was next conjugated with Brilliant Violet 421-streptavidin (Biolegend) at a molar ratio of 4:1. Splenic MNCs were incubated with HBsAg probes for 20 minutes, and then stained with Live/Dead Fixable Blue Dead Cells Stain Kit (Thermofisher) for 5 minutes. After washing, cells were incubated with antibody cocktails for 20 minutes at 4 ℃ in dark. Flow cytometry analysis was carried out on BD FACSymphony A3 (BD Biosciences). Data was analyzed using FlowJo V10.8 software (Tree Star). Fluorochrome-conjugated antibodies used in this study are listed in Supplemental Table 2.

**Evaluation of antigen-specific T cell response**

3 million splenic MNCs were seeded per well into 96-well U-bottom plates and incubated with HBsAg overlapping peptides (15-mers overlapping by 10 amino acids, GenScript Probio) at a concentration of 10μg/mL for 16 hours in the presence of 5μg/mL brefeldin A (Biolegend). Splenic MNCs stimulated with 2μg/mL staphylococcal enterotoxin B (SEB, Creative Diagnostics) were used as positive control. Cytokine production of antigen-specific T cells was evaluated by surface and intracellular staining using Fixation/Permeabilization Solution Kit (BD Biosciences) according to the manuals. Frequencies of cytokine-producing T cells were determined by FACS analysis. Background cytokine staining was subtracted, as defined by staining in the samples incubated with medium alone. Flow cytometry analysis was carried out on BD FACSymphony A3 (BD Biosciences). Data was analyzed using FlowJo V10.8 software (Tree Star). Fluorochrome-conjugated antibodies used in this study are listed in Supplemental Table 2.

**Supplemental Figure 1**

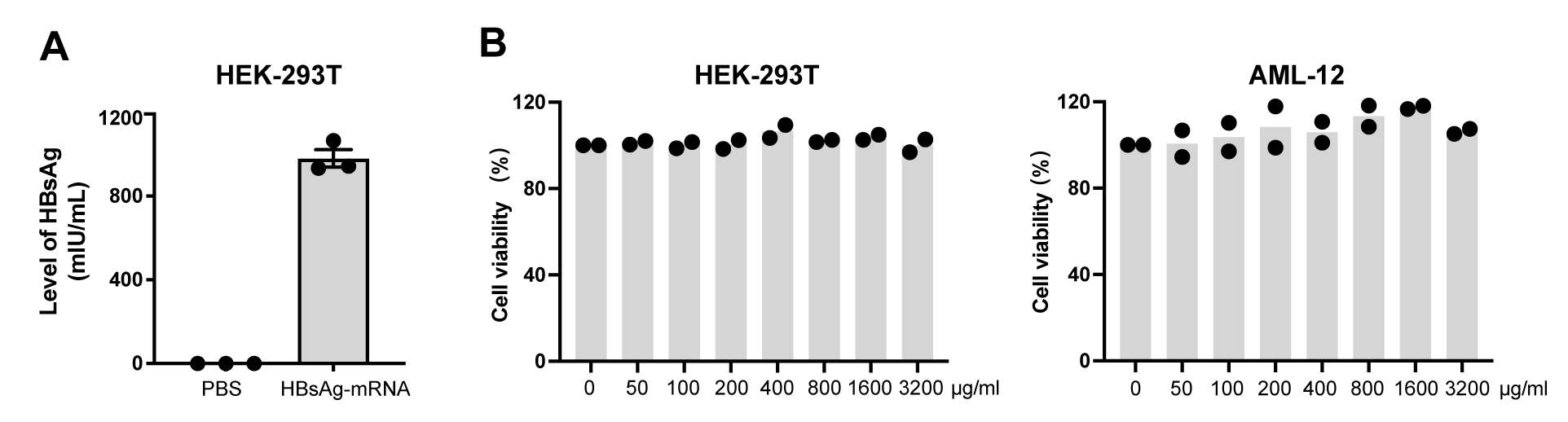

**Supplemental Figure 1. Translation of HBsAg-encoding mRNA and cytotoxicity of HBV mRNA vaccine.**

(A) HEK-293T cells transfected with 2μg m1Ψ-modified HBsAg-encoding mRNAs were harvested 48 hours after transfection. HBsAg levels in cell lysates were detected. Data compiled from three independent experiments are shown as mean ± SEM. (B) HEK-293T and AML-12 cells were treated with escalating doses of HBV mRNA vaccines for 24 hours. Cell viability was determined with CCK8 assays and depicted as percentage relative to untreated cells. Data from two independent experiments are shown.

**Supplemental Figure 2**

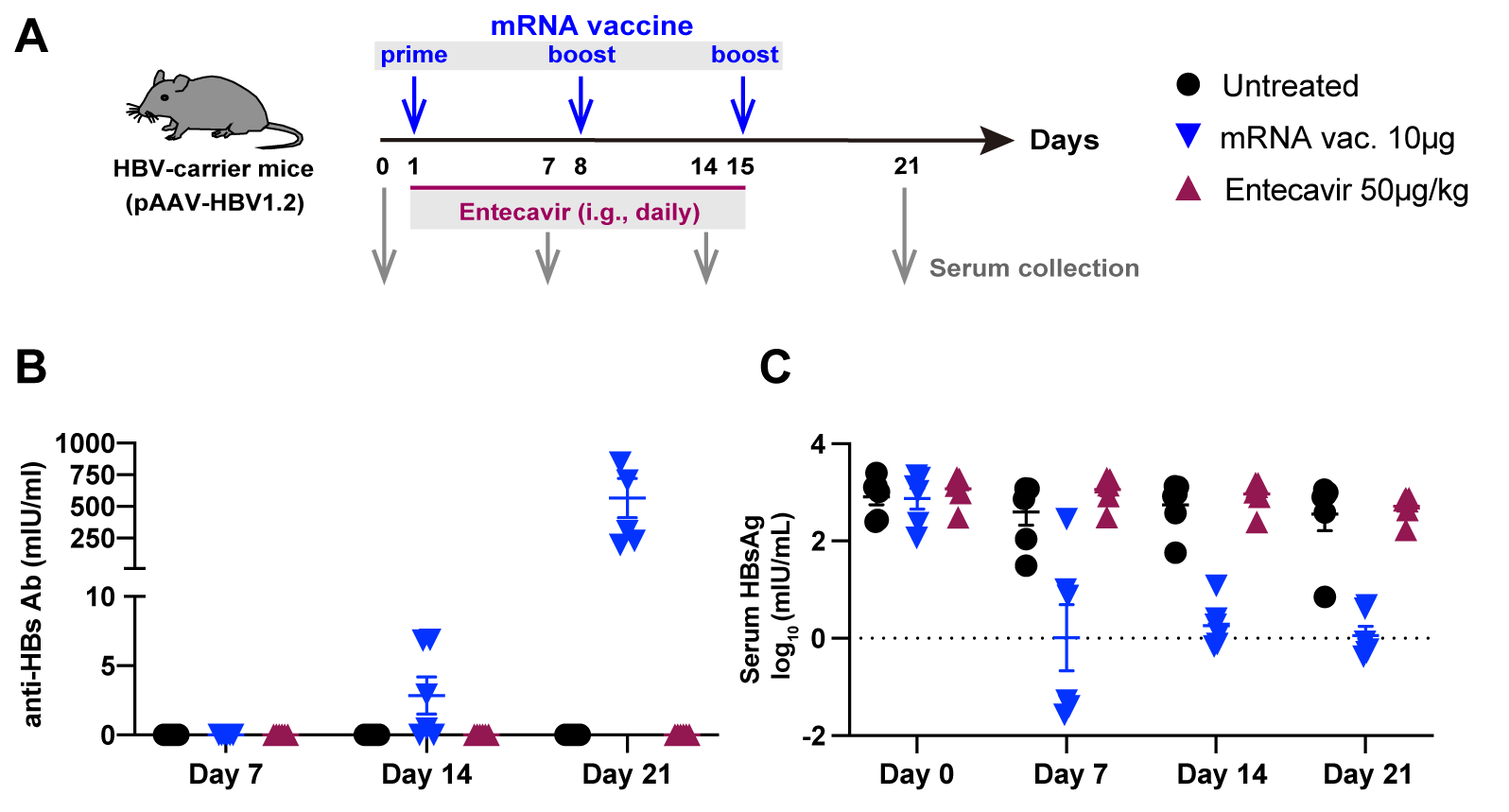

**Supplemental Figure 2. HBV mRNA vaccine showed superior efficacy over Entecavir.**

(A) pAAV/HBV1.2-transduced HBV-carrier mice (n=6/group) were either immunized i.m. with 10μg HBV mRNA vaccines three times at a 1-week interval or were orally treated with Entecavir (ETV, 50μg/kg) for 15 days consecutively. HBV-carrier mice administered with PBS were used as control. Sera samples were collected at the indicated time points. (B) Levels of anti-HBs Abs were evaluated. (C) Serum HBsAg were measured by chemiluminescence immunoassay (CLIA). Data are shown as Mean ± SEM.

**Supplemental Figure 3**

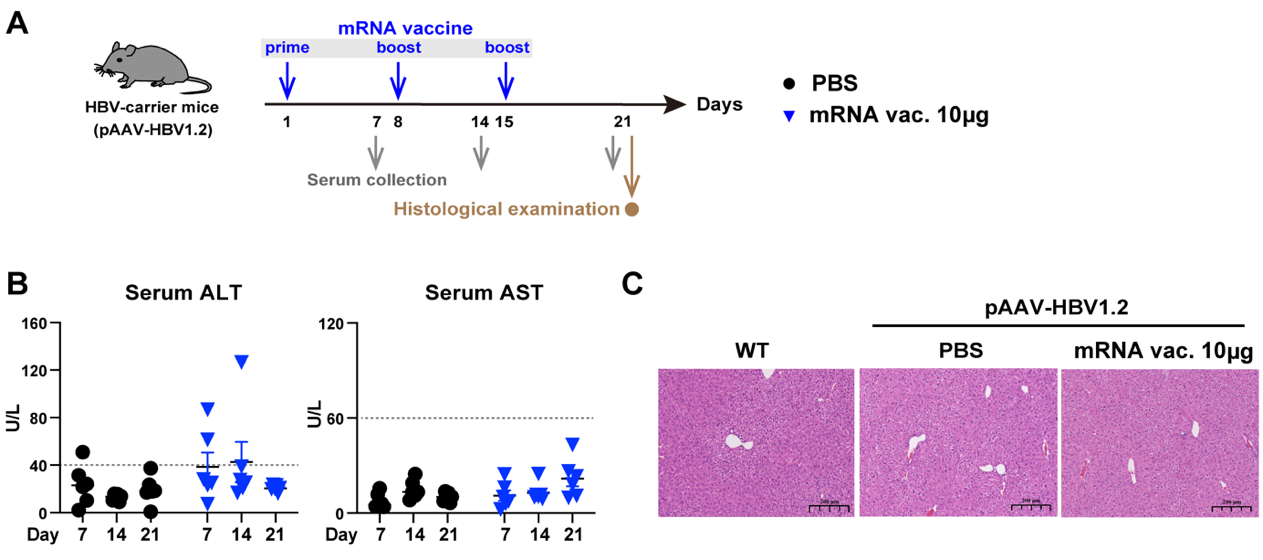

**Supplemental Figure 3. HBV mRNA vaccine induced very limited hepatotoxicity and liver injury.**

(A) pAAV/HBV1.2-transduced HBV-carrier mice (n=6/group) were immunized i.m. with 10μg HBV mRNA vaccines three times at a 1-week interval. Sera samples and liver tissues were collected at the indicated time points. (B) Levels of serum ALT and AST were assessed. Data are shown as Mean ± SEM. The dotted line depicts reported normal levels of the two parameters. (C) Hematoxylin and eosin (H&E) staining for histopathologic analysis of liver tissues collected at the indicated time point. Scale bar: 200 µm; WT: wide-type healthy mice.

**Supplemental Figure 4**

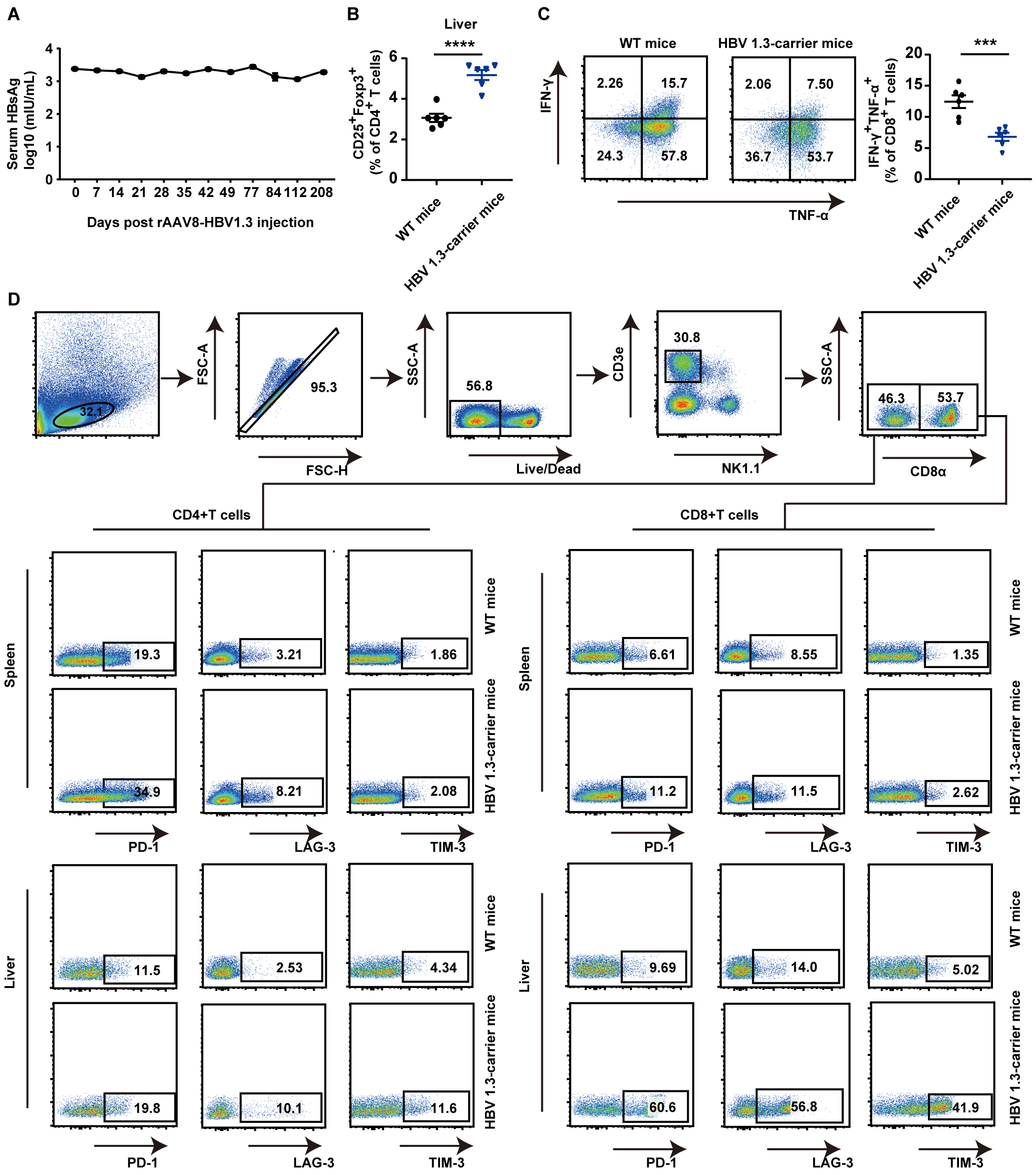

**Supplemental Figure 4. rAAV8-HBV1.3-transduced mice showed persistent HBV viremia and immunotolerance.**

(A) C57BL/6J mice (n=6) were intravenously injected with 1 × 10^10^ vector genome equivalent of rAAV8-HBV1.3. Serum HBsAg levels were detected at the indicated time points after rAAV8-HBV1.3 injection. (B) Frequencies of hepatic CD4^+^CD25^+^Foxp3^+^ cells from rAAV8-HBV1.3 mice and WT mice are shown. (C) Hepatic MNCs (2 × 10^6^) from rAAV8-HBV1.3 mice and WT mice were stimulated with PMA/ionomycin in vitro for 4 hours in the presence of brefeldin A (5μg/mL), and the production of IFN-γ and TNF-α by CD8^+^ T cells was analyzed by flow cytometry. (D) Expression of PD-1, LAG-3, and TIM-3 on hepatic and splenic CD4^+^ and CD8^+^ T cells from rAAV8-HBV1.3 mice and WT mice were analyzed by flow cytometry. Mean fluorescence intensity (MFI) values are shown. ****p* ≤ 0.001, *****p* ≤ 0.0001.

**Supplemental Figure 5**

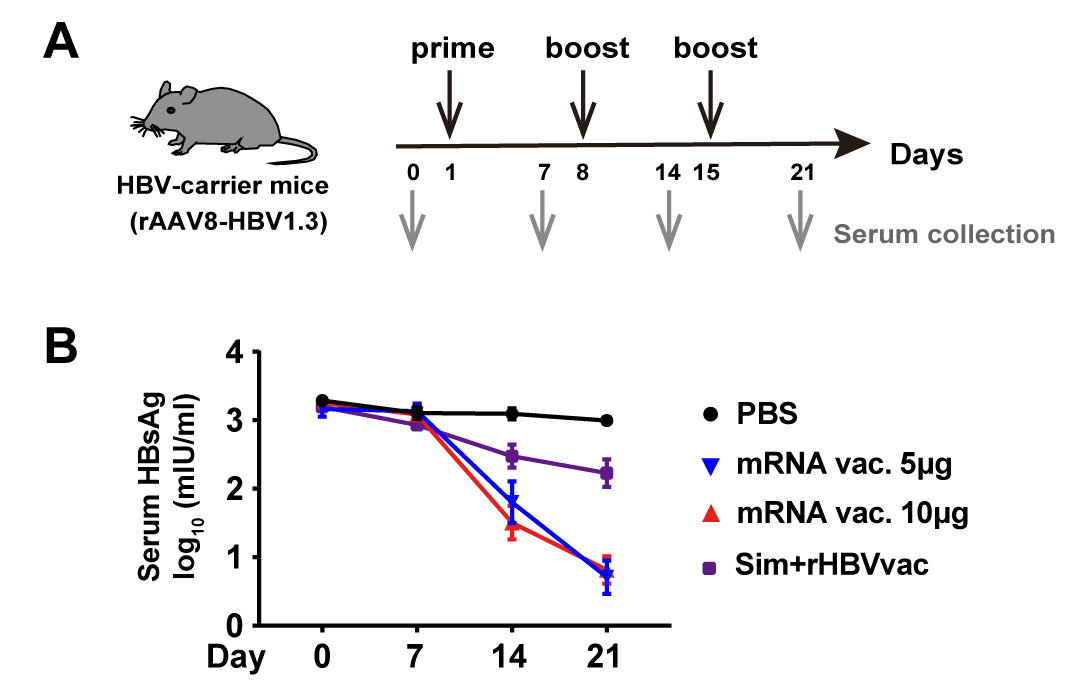

**Supplemental Figure 5. HBV mRNA vaccine showed superior efficacy over recombinant therapeutic vaccine in viral clearance.**

(A) rAAV/HBV1.3-transduced HBV-carrier mice (n=6/group) were immunized i.m. with HBV mRNA vaccines or recombinant HBV therapeutic vaccines (“Sim+rHBV”) three times at a 1-week interval. HBV-carrier mice administered with PBS were used as control. Sera samples were collected at the indicated time points. (B) Serum HBsAg levels were detected by chemiluminescence immunoassay (CLIA). Data are shown as Mean ± SEM.

**Supplemental Figure 6**

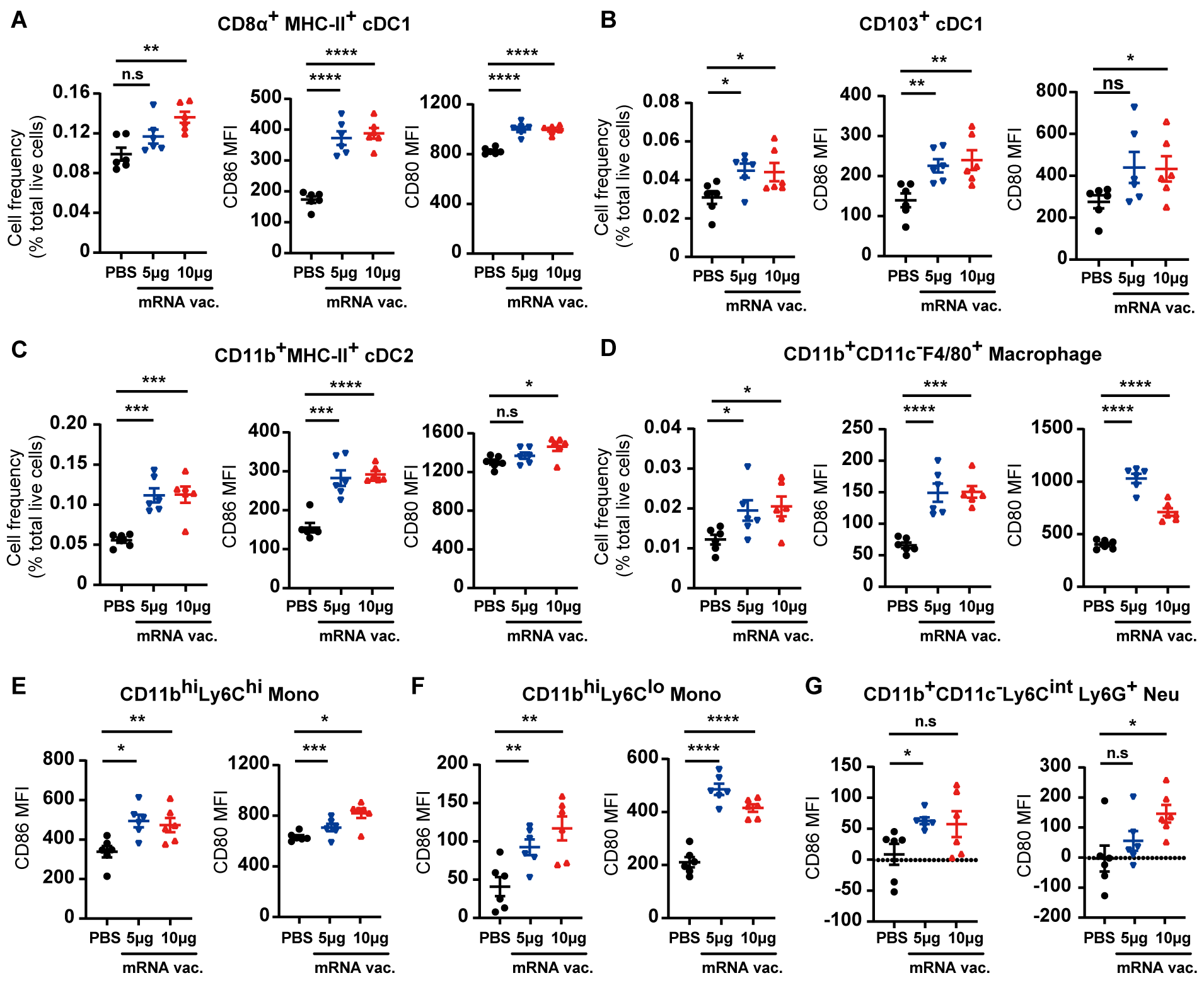

**Supplemental Figure 6. HBV mRNA vaccine induced strong innate immune activation in HBV-carrier mice.**

rAAV/HBV1.3-transduced HBV-carrier mice (n=6/group) were i.m. injected with 5μg or 10μg HBV mRNA vaccine. Splenic mononuclear cells (MNCs) were collected 12 hours after immunization. (A-D) Frequencies of CD8α^+^ cDC1s, CD103^+^ cDC1s, CD11b^+^ cDC2s, and F4/80^+^ Macrophages and expression of CD80 and CD86 on these cell subsets were evaluated by flow cytometry. Mean fluorescence intensity (MFI) values are shown. (E-G) Expression of CD80 and CD86 on CD11b^hi^Ly6C^lo^ monocytes, CD11b^hi^Ly6C^hi^ monocytes and CD11b^+^ CD11c^-^ Ly6C^int^ Ly6G^+^ neutrophils were analyzed by flow cytometry. MFI values are shown. **p* ≤ 0.05, ***p* ≤ 0.01, ****p* ≤ 0.001, *****p* ≤ 0.0001.

**Supplemental Figure 7**

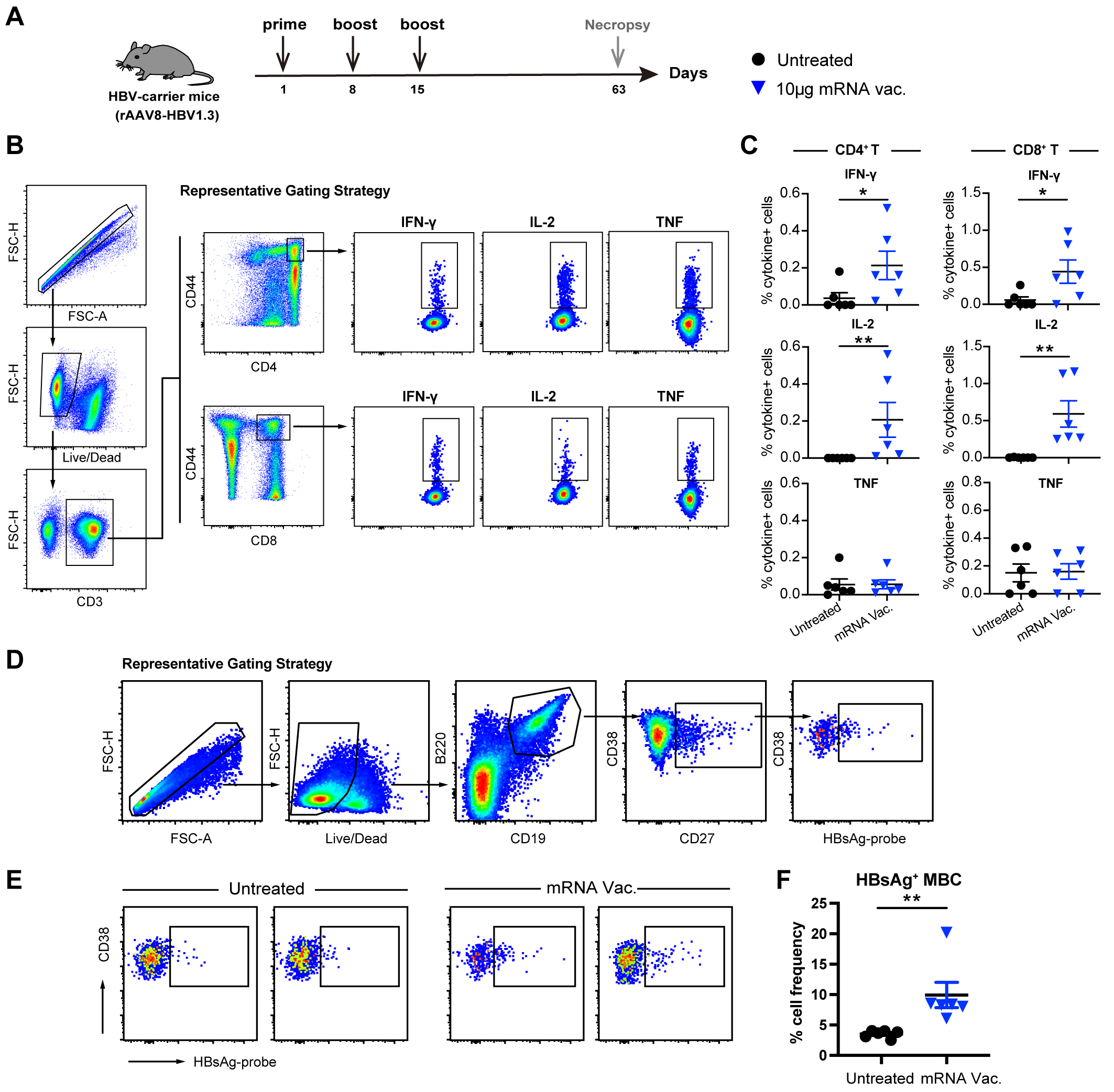

**Supplemental Figure 7. HBV mRNA vaccine induced robust HBsAg-specific Th1-biased T cell and memory B cell responses.**

(A) rAAV/HBV1.3-transduced HBV-carrier mice (n=6/group) were immunized i.m. with 10μg HBV mRNA vaccines three times at a 1-week interval. HBV-carrier mice administered with PBS were used as control. Spleens were collected 48 days after the 3^rd^ vaccine dose. (B-C) Splenic cells were stimulated with 10μg/ml HBsAg overlapping peptides (15-mers overlapping by 10 amino acids) for 16 hours in the presence of brefeldin A. Cells stimulated with staphylococcal Enterotoxin B (SEB) were used as positive control. (B) Representative gating of SEB-stimulated cells secreting cytokines. (C) Quantification of cytokine-producing antigen-specific CD4^+^ and CD8^+^ T cells following overlapping peptides stimulation. (D-F) Frequencies of HBsAg-specific memory B cells (MBCs) in spleens were evaluated. (D) Representative gating of HBsAg-specific MBCs. (E) Data from two representative animals from each group are shown. (F) Frequencies of HBsAg-specific MBCs in spleens of mice from the indicated groups. Data are shown as Mean ± SEM. *p ≤ 0.05, **p ≤ 0.01.

**Supplemental Tables**

**Table S1. List of primers for HBV DNA and RNA quantification.**

| **REAGENT** | **Primers** |
| --- | --- |
| mouse-GAPDH-F | 5’-AGG TCG GTG TGA ACG GAT TTG-3’ |
| mouse-GAPDH-R | 5’-TGT AGA CCA TGT AGT TGA GGT CA-3’ |
| HBV-cccDNA-F | 5’-CGT CTG TGC CTT CTC ATC TGC-3’ |
| HBV-cccDNA-R | 5’-GCA CAG CTT GGA GGC TTG AA-3’ |
| HBV-DNA-F | 5’-CAC ATC AGG ATT CCT AGG ACC-3’ |
| HBV-DNA-R | 5’-GGT GAGTGA TTG GAG GTT G-3’ |
| HBV-total-RNA-F | 5’-TCA CCA GCA CCA TGC AAC-3’ |
| HBV-total-RNA-R | 5’-AAG CCA CCC AAG GCA CAG-3 |
| HBV-3.5kb-RNA-F | 5’-GAG TGT GGA TTC GCACTC C-3’ |
| HBV-3.5kb-RNA-R | 5’-GAG GCG AGG GAG TTC TTC T-3’ |

**Table S2. List of anti-mouse antibodies used for FACS analysis.**

| **Antibody** | **Clone** | **Manufacturer** |
| --- | --- | --- |
| CD45R/B220 | RA3-6B2 | Biolegend |
| CD19 | 6D5 | Biolegend |
| CD38 | 90 | Biolegend |
| CD27 | LG.3A10 | Biolegend |
| CD3 | 17A2 | Biolegend |
| CD4 | GK1.5 | Biolegend |
| CD44 | IM7 | Biolegend |
| IFN-γ | XMG1.2 | Biolegend |
| TNF | MP6-XT22 | Biolegend |
| IL-2 | JES6-5H4 | Biolegend |
| PD-1 | 29F.1A12 | Biolegend |
| LAG-3 | C9B7W | Biolegend |
| TIM-3 | RMT3-23 | Biolegend |
| CD103 | 2E7 | Biolegend |
| CD11c | N418 | Biolegend |
| MHC-Ⅱ | M5/114.15.2 | Biolegend |
| CD80 | 16-10A1 | Biolegend |
| CD86 | IT2.2 | Biolegend |
| Gr-1 | RB6-8C5 | eBioscience |
| Ly6C | HK1.4 | eBioscience |
| CD11b | M1/70 | eBioscience |
| CD25 | PC61.5 | eBioscience |
| Foxp3 | FJK-16s | eBioscience |
| F4/80 | BM8 | eBioscience |
| CD8α | 53-6.7 | BD |
